# Discovery of CD28-Targeted Small Molecule Inhibitors of T Cell Co-stimulation Using Affinity Selection-Mass Spectrometry (AS-MS) and Ex Vivo Validation

**DOI:** 10.1101/2025.07.31.667814

**Authors:** Saurabh Upadhyay, Sungwoo Cho, Hossam Nada, Moustafa T. Gabr

## Abstract

CD28 is a key T cell co-stimulatory receptor implicated in antitumor immunity and immune-related disorders, yet no small molecule modulators of CD28 have reached clinical development. Here, we report the discovery and characterization of small molecule CD28 antagonists identified through affinity selection-mass spectrometry (AS-MS). Subsequent catalog-based structure–activity relationship (SAR) optimization led to the identification of two lead compounds, **5MS-5** and **19MS-5**, which exhibit direct CD28 binding and potent inhibition of CD28–B7 interactions in cellular reporter assays. In vitro pharmacokinetic profiling demonstrated favorable solubility, metabolic stability, and permeability, alongside low off-target liabilities. Functionally, both compounds suppressed cytokine production in primary human T cells co-cultured with tumor spheroids and human epithelial tissues, validating their ability to inhibit CD28-driven immune activation in physiologically relevant models. These findings establish **5MS-5** and **19MS-5** as promising CD28 inhibitors and provide a foundation for developing orally bioavailable immunomodulators targeting T cell co-stimulation.

**Table of Contents artwork:** 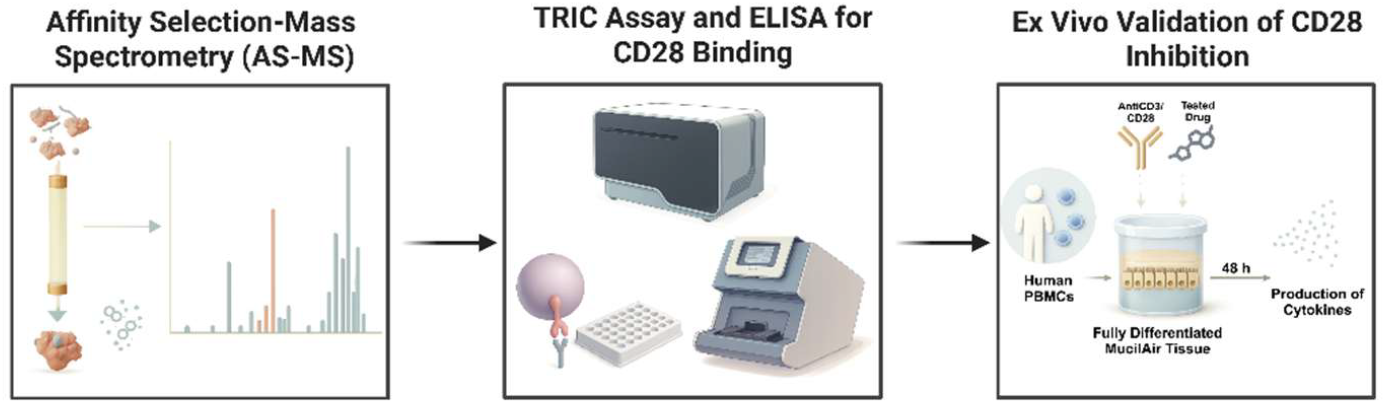

## INTRODUCTION

Immune checkpoint receptors regulate the initiation, amplitude, and resolution of T cell responses.^1–4^ Among these, Cluster of Differentiation 28 (CD28) is the primary co-stimulatory receptor on naïve T cells, delivering a co-stimulatory signal upon engagement with CD80 (B7-1) or CD86 (B7-2) on antigen-presenting cells.^5, 6^ This interaction potentiates T cell receptor (TCR) signaling, promoting IL-2 production and T cell expansion.^7–9^ However, dysregulated CD28 signaling contributes to autoimmune pathology, transplant rejection, and tumor immune evasion.^3,10–12^ Despite its central role, CD28 remains largely untargeted by small molecules.^13, 14^ CD28 lacks a catalytic site and presents a flexible, shallow protein–protein interaction (PPI) interface that is challenging to target using small molecules.^15, 16^ Moreover, therapeutic targeting of CD28 requires exceptional precision, as evidenced by the severe cytokine release syndrome observed in the TGN1412 clinical trial.^17^

Several biologics targeting CD28 have been developed, including FR104 (a monovalent anti-CD28 Fab’), lulizumab pegol, and BMS-931699, which aim to suppress T cell costimulation in autoimmunity and transplantation.^18–20^ While promising, these agents face limitations including poor tissue penetration, parenteral delivery, immunogenicity, and limited pharmacokinetic (PK) tunability.^21^ Next-generation approaches like CD28-targeted CAR-T cells and bispecific antibodies offer mechanistic sophistication but are costly, complex to manufacture, and difficult to dose-adjust.^22, 23^

These limitations underscore the unmet need for chemically defined, reversible, and tunable small molecule CD28 modulators that can be readily integrated into combination immunotherapy regimens. Notably, resistance to current immune checkpoint inhibitors, particularly PD-1 and CTLA-4 blockade, is increasingly linked to sustained CD28 costimulatory signaling.^24^ Even with PD-1 blockade, CD28 engagement with CD80/CD86 can restore T cell activation promoting tumor immune escape and therapeutic failure.^25^ Emerging data from both preclinical and clinical studies highlight CD28 as a key driver of resistance, and its co-targeting has been shown to enhance antitumor responses in resistant settings.^3^ Thus, CD28 is a mechanistically distinct target for addressing resistance to checkpoint blockade.

To address this therapeutic challenge, we employed affinity selection mass spectrometry (AS-MS), a high-throughput, label-free biophysical platform that enables detection of protein–ligand interactions in solution with high specificity.^26, 27^ Unlike conventional biochemical high-throughput screening (HTS), which relies on activity-coupled or fluorescence-based readouts, AS-MS facilitates direct, function-independent identification of ligands that bind target proteins.^28^ However, applying AS-MS to a structurally dynamic, non-enzymatic target such as CD28 required substantial methodological innovation. We established a size exclusion chromatography (SEC)–coupled AS-MS pipeline, optimizing protein and buffer conditions, library complexity, and SEC resolution to enable selective isolation of CD28–ligand complexes and subsequent high-resolution mass spectrometric identification of retained binders. Hits identified from the screen were advanced through a tiered validation cascade, including CD28 binding studies and functional assays assessing CD28:B7-1 inhibition and downstream co-stimulation blockade. This strategy yielded small molecules that directly bind CD28 and disrupt its interaction with B7 ligands, demonstrating the feasibility of chemically modulating this historically intractable checkpoint. Guided by structure-activity relationship (SAR) by catalog studies, we further optimized functional potency, resulting in tractable chemical scaffolds suitable for medicinal chemistry refinement. This framework draws on non-canonical, structure-guided strategies used to target PKM2, DAAO, and GPCRs.^29–31^

Here, we report the discovery and optimization of potent small molecule CD28 inhibitors that directly bind CD28 and disrupt its interaction with B7 ligands. These compounds represent a significant advance in targeting co-stimulatory pathways using chemically defined scaffolds with favorable preliminary PK profiles. We demonstrate their translational relevance using ex vivo co-culture systems comprising peripheral blood mononuclear cells (PBMCs) and either human tumor or mucosal tissues, capturing clinically relevant immune–tissue interactions. By selectively and reversibly modulating CD28 signaling, these small molecules offer a tractable alternative to antibody-based agents, with advantages including tunable PK, enhanced tissue penetration, and reduced immunogenicity. This study builds on our lab’s broader efforts to develop small molecule modulators of immune checkpoints^32–40^ and establishes a framework for CD28-targeted immunotherapies with broad applicability across oncology, autoimmunity, and transplantation—including in settings of resistance to current checkpoint blockade.

## RESULTS AND DISCUSSION

### High-Throughput Mass Spectrometry-Based Screening

To identify small molecule ligands capable of binding CD28, we developed a two-dimensional size-exclusion chromatography–coupled affinity selection mass spectrometry (SEC-coupled AS-MS) platform optimized for native, solution-phase interactions (Figure 1A). CD28 protein was incubated with pooled small molecule libraries and subjected to SEC to isolate protein–ligand complexes, which were then denatured and analyzed by high-resolution LC-MS to detect co-eluting ligands.

**Figure 1.**
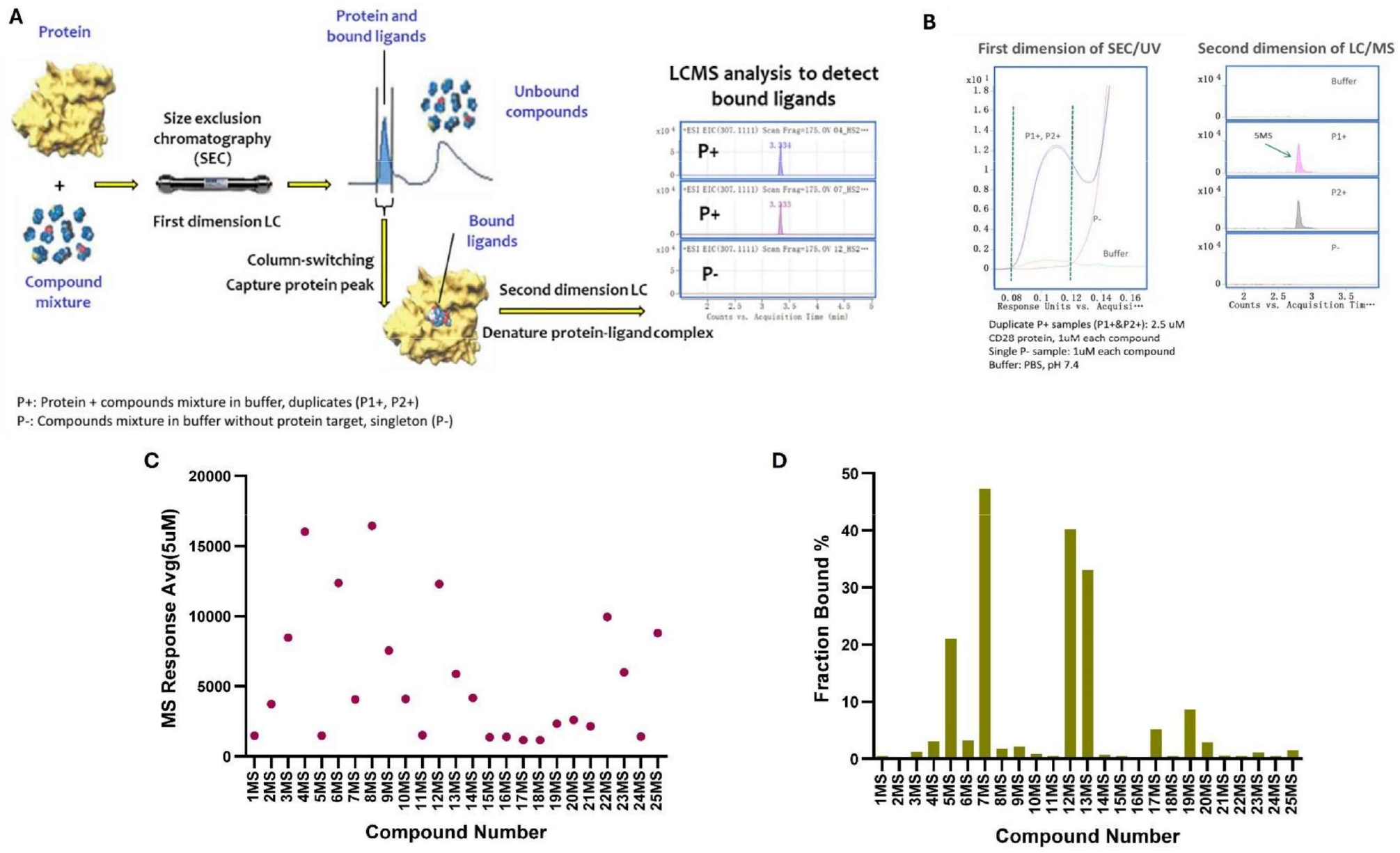
Two-dimensional SEC-coupled AS-MS screening identifies CD28-targeted small molecules. **(A)** Schematic of the SEC-coupled AS-MS workflow. A total of 6,336 compounds were screened in pooled format using CD28 protein, followed by SEC and LC-MS to detect protein-bound ligands. **(B)** Representative SEC/UV chromatograms from the primary screen using 2.5 μM CD28 and 1 μM compound concentration. A distinct protein peak is observed in CD28 + compound samples (P1+, P2+) but absent in the compound-only control (P−), confirming successful protein separation. **(C)** Hit selection was based on mass error ≤5 ppm, absence in P− control, P+/P− intensity ratio ≥3, consistent retention time, clear isotopic pattern, and signal >1000 counts. A total of 56 compounds were initially selected, and 25 were confirmed as CD28 binders. **(D)** Fraction bound (%) of selected compounds from the retest screen using 5 μM CD28. Compounds **5MS, 7MS, 12MS, 13MS, 17MS, 19MS**, and **20MS** showed the highest Fb% values (>30%), indicating strong binding to CD28.

In the primary screen, ~6,336 compounds (from Enamine Diversity Chemical Library) were evaluated at 1 μM ligand concentration using 2.5 μM CD28, with compound pools containing ~1,050 molecules each. SEC/UV chromatograms revealed distinct protein peaks in CD28-containing samples (P1+ and P2+), which were absent in compound-only controls (P−), confirming successful separation of protein–ligand complexes from unbound compounds (Figure 1B). LC-MS analysis identified 52 preliminary binders based on stringent criteria, including mass error ≤5 ppm, absence in the P− control, P+/P− intensity ratio ≥3, consistent retention time, clean isotopic patterns, and signal intensity >1,000 counts.

To confirm specificity, these 52 candidates were retested using 5 μM CD28 concentration and smaller pools (~10 compounds per pool). This secondary screen yielded 25 reproducible hits (0.39% hit rate) that met the same rigorous thresholds. The chemical structures of these 25 compounds are shown in Table S1. As shown in Figure 1C, the hits exhibited variable MS signal intensities, with some compounds—such as **4MS, 5MS, 7MS**, and **12MS**—showing particularly remarkable responses. Binding strength was further assessed by calculating the fraction bound (Fb%), which quantifies the proportion of each ligand retained in the protein peak. Figure 1D highlights compounds **5MS, 7MS, 12MS, 13MS** and **19MS** as top binders, with Fb% values of 20%, 47%, 44%, 33% and 9%, respectively. Together, these quantitative metrics validate the specificity and robustness of the CD28–ligand interactions identified, laying a strong foundation to evaluate all 25 compounds in downstream biophysical and functional studies. The prioritized hits represent chemically tractable starting points for further validation and structure-based optimization targeting CD28-mediated T cell co-stimulation.

### Hit Validation

Dianthus is an instrument that quantifies molecular interactions by measuring fluorescence intensity changes triggered by temperature shifts, a principle known as temperature-related intensity change (TRIC). To validate the binding of compounds identified by SEC-coupled AS-MS, all 25 confirmed hits were evaluated using a single-dose fluorescence-based binding screen on the Dianthus platform (Figure 2A). Compounds were tested at 100 μM in PBST buffer (PBS with 0.05% Tween-20) supplemented with 2% DMSO, and binding to CD28 was quantified based on changes in fluorescence signal relative to a buffer-only control. Compounds were classified as binders if their signal deviated by at least threefold from the background variation. Based on this criterion, three compounds (**5MS, 19MS**, and **20MS**) exhibited significant signal shifts indicative of specific binding and were advanced for further analysis (Figure 2A).

**Figure 2.**
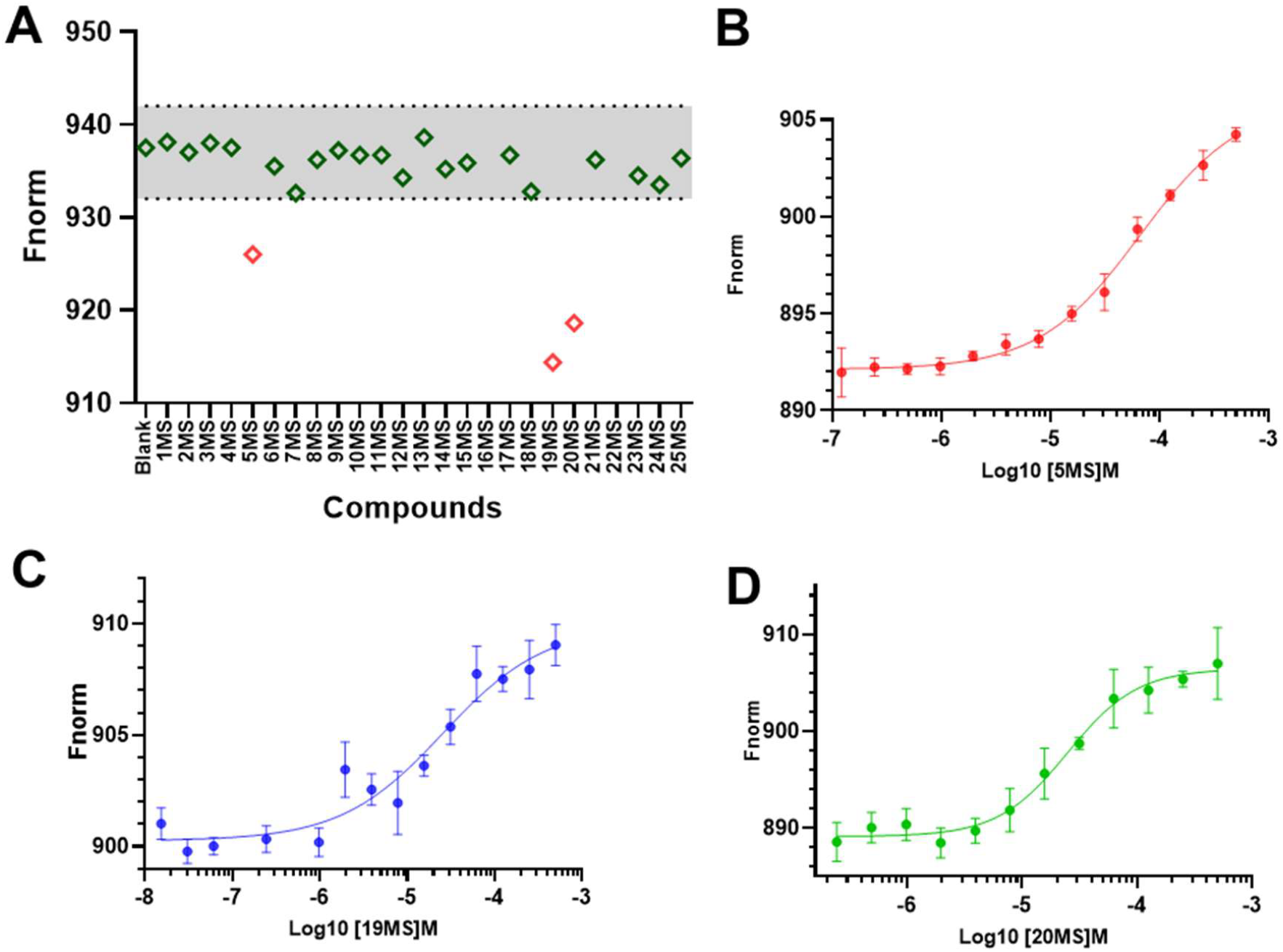
Binding validation of CD28 ligands using Dianthus and MST. **(A)** Single-dose binding responses of 25 hits at 100 μM using the Dianthus platform in PBST buffer (PBS + 0.05% Tween-20) with 2% DMSO. Compounds higher or lower three times the SD of CD28 only (Blank) were selected for dose dependent analysis. **(B–D)** Dose–response MST binding curves for compounds **5MS, 19MS**, and **20MS**, respectively. Each compound was prepared at an initial concentration of 500 μM and subjected to a 14-point 1:1 serial dilution (final range: 500 to 0.06 μM). Data represent mean ± standard error mean (SEM) from n = 5 independent measurements.

To determine the binding affinity of these three compounds, we performed dose–response measurements using microscale thermophoresis (MST). Although both Dianthus and MST (Monolith) are TRIC-based platforms, MST offers higher sensitivity, reduced sample volume requirements, and finer control over thermal gradients. This sequential use of Dianthus for primary screening and Monolith for quantitative follow-up represents a standard and recommended workflow for TRIC-based hit validation. Compounds were serially diluted (500 μM to 0.06 μM, 1:1 dilution) in PBST with 2% DMSO, and incubated with a constant concentration of CD28 protein. The resulting thermophoresis signals were fitted to generate binding curves and calculate equilibrium dissociation constants (K_d_). All three compounds displayed clear, concentration-dependent binding to CD28. The determined K_d_ values were: **5MS** (50.32 ± 13.14 μM, Figure 2B), **19MS** (18.25 ± 5.40 μM, Figure 2C), and **20MS** (24.28 ± 11.80 μM, Figure 2D). These results confirm the direct and specific binding of these small molecules to CD28 under physiologically relevant buffer conditions. Their validated affinity profiles provide a strong rationale for initiating SAR studies to optimize potency.

### Preliminary SAR-by-Catalog Approach

To improve the potency of the initial CD28 hits, we conducted a SAR-by-catalog study using the three validated hit compounds (**5MS, 19MS**, and **20MS**) as starting points (Figure 3). Twenty structurally related analogs were sourced from the Enamine database based on scaffold similarity (Figure 3). For the **5MS** series (Figure 3A), modifications were introduced at the aryl and heteroaryl regions, including methoxy (**5MS-1, 5MS-2, 5MS-4, 5MS-5**, and **5MS-6**), acetyl (**5MS-3**), halogen (**5MS-5, 5MS-8**, and **5MS-9**), and fused ring substitutions (**5MS-7**). The **19MS** analogs (Figure 3B) explored variations in the aryl moiety and linker region. For example, **19MS-1** and **19MS-2** feature phenoxy and pyridyl moieties, respectively. **19MS-6** and **19MS-9** incorporate diverse ring systems such as indole and triazole, respectively. Three analogs (**19MS-5, 19MS-7**, and **19MS-9**) feature substituted pyrazole rings. In contrast, the **20MS** analogs (Figure 3C) incorporated subtle variations to the linker and the incorporation of heterocycle rings, such as tetrahydropyran (**20MS-1, 20MS-2**).

**Figure 3.**
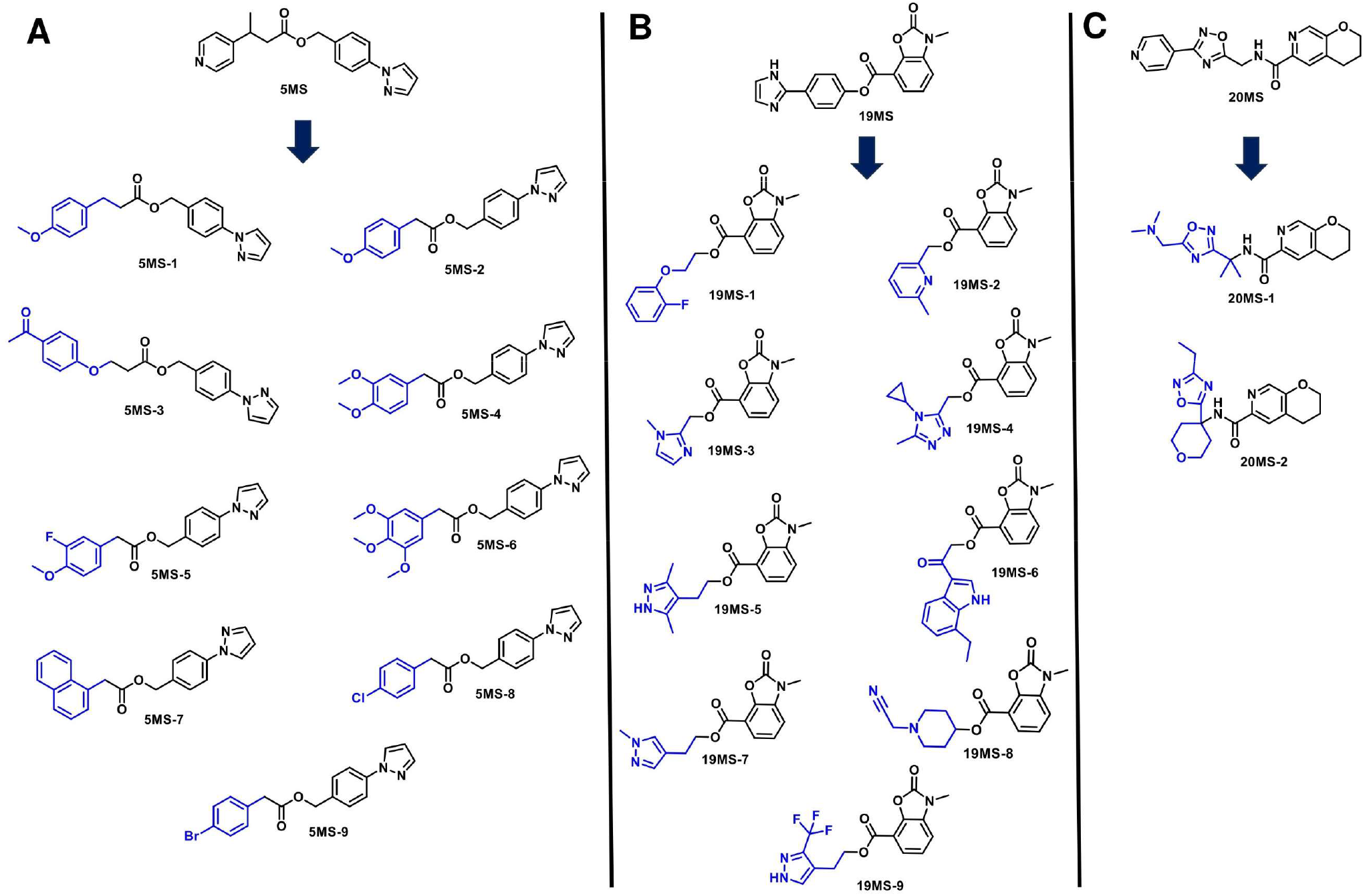
SAR-by-catalog approach for CD28-binding compounds. **(A)** Analog series derived from hit compound **5MS**, highlighting substitutions on the aromatic and heterocyclic moieties. **(B)** Analog series based on compound **19MS**, exploring variations in the aromatic moieties and linkers. **(C)** Analog series stemming from **20MS** with modifications to the side chain. Analog selection was guided by scaffold similarity and commercial availability (Enamine database). Modified regions are highlighted in blue.

The 20 catalog analogs were initially screened for CD28 binding using the TRIC-based Dianthus platform (Figure 4). Each compound was tested at 100 μM in PBST buffer (PBS containing 0.05% Tween-20) supplemented with 2% DMSO. Fluorescence signal changes were compared to a buffer-only negative control, and compounds were classified as binders if their signal deviated by at least threefold from baseline variation. This primary screen (Figure 4) identified three potential binders, one analog from the **5MS** series (**5MS-5**) and two from the **19MS** series (**19MS-2** and **19MS-5**).

**Figure 4.**
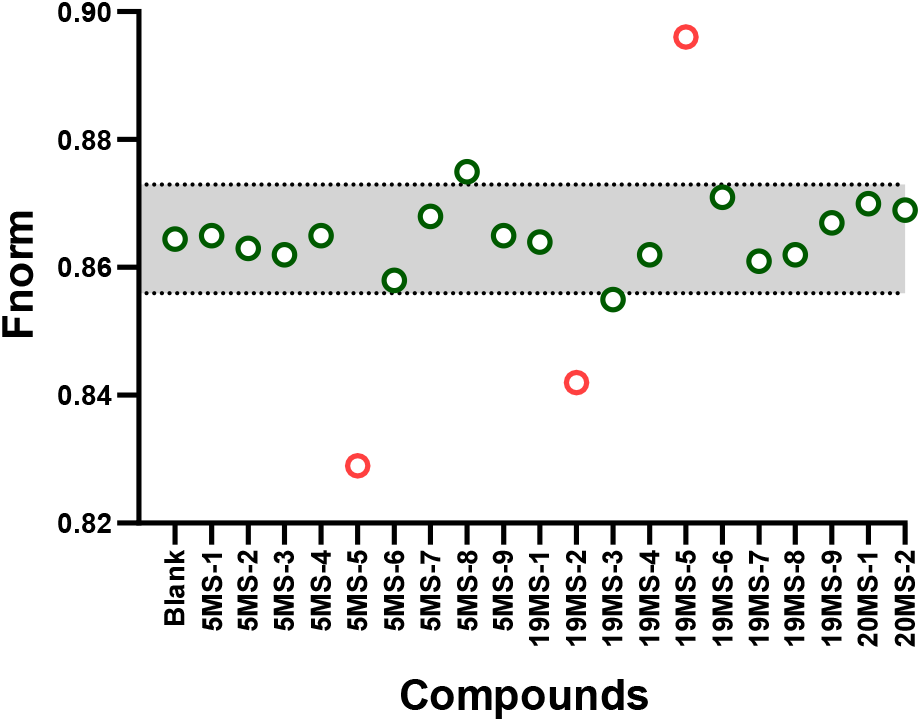
SAR-driven analog screening and binding validation of CD28 ligands. Single-dose binding screen of 20 structural analogs tested at 100 μM using the Dianthus system in PBST buffer (PBS + 0.05% Tween-20) with 2% DMSO. Fluorescence shifts were compared to a buffer-only control. Compounds showing signal changes greater than three times the standard deviation of the control were considered binders. One **5MS** analog and two **19MS** analogs met this criterion.

To validate these optimized compounds and quantify their binding affinities, all three compounds were subjected to dose–response analysis using MST. Compounds were serially diluted 1:1 from 500 μM to 0.06 μM in PBST with 2% DMSO, while the concentration of CD28 protein was kept constant. Among the tested analogs, **5MS-5** and **19MS-5** exhibited clear, concentration-dependent binding curves with K_d_ values of 24.13 ± 6.28 μM and 12.48 ± 3.69 μM, respectively (Figure 5). Structurally, **5MS-5** features *meta*-fluoro and *para*-methoxy substitutions on the aryl ring, which likely enhances favorable interactions within the CD28 binding pocket. Similarly, the pyrazole analog **19MS-5** retains the core heterocyclic lactam motif of **19MS** while introducing a flexible linker that may improve binding through additional interactions or conformational adaptability. In contrast, the remaining analogs, including those from the **20MS** series, did not show measurable CD28 binding, suggesting a limited SAR window for this scaffold. These results demonstrate that SAR-by-catalog screening successfully identified analogs with improved CD28 binding affinity, particularly for the **5MS** and **19MS** scaffolds. The enhanced potency of **5MS-5** and **19MS-5** highlights their promise as lead compounds for further optimization.

**Figure 5.**
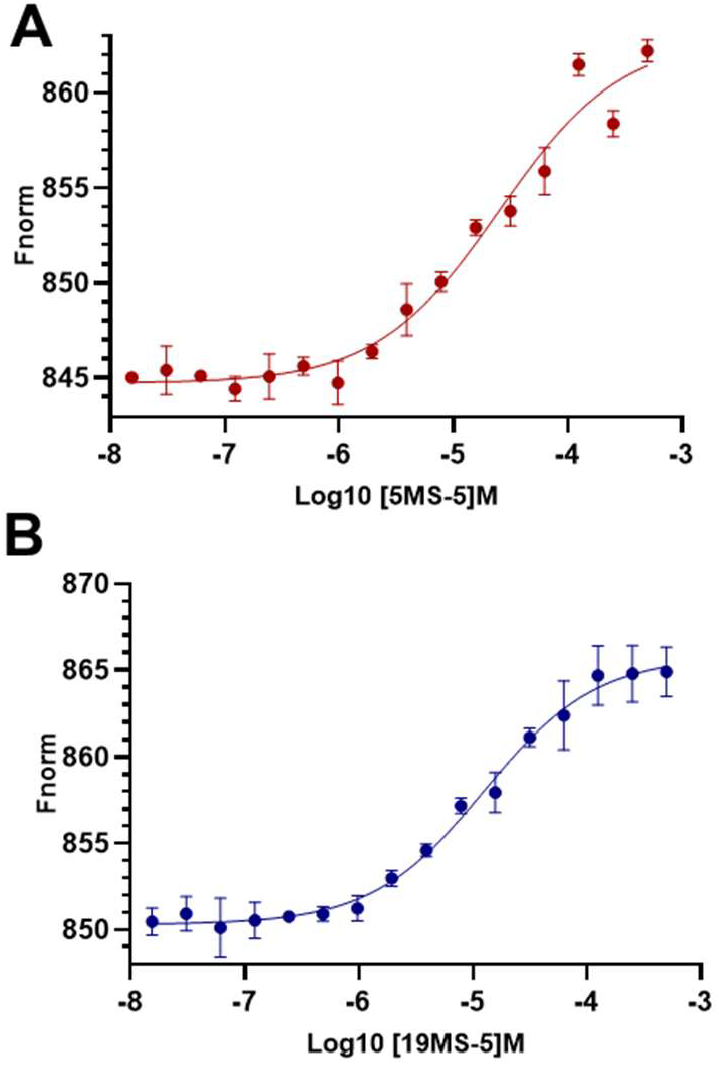
MST dose–response curves for validated analogs 5MS-5 (A) and 19MS-5 (B). Each compound was serially diluted from 500 to 0.06 μM in PBST with 2% DMSO, while CD28 protein concentration was held constant. KD values were determined by nonlinear regression using GraphPad Prism, fitted to a 1:1 binding model. Data are presented as mean ± SEM from n = 5 independent experiments.

### Inhibition of CD28 Interactions

Following biophysical validation of CD28 binding, we next evaluated the ability of the optimized compounds (**5MS-5** and **19MS-5**) to functionally inhibit the CD28:B7-1 interaction. This protein– protein interaction is essential for T cell co-stimulation, making it a critical target for checkpoint modulation. To this end, an ELISA-based inhibition assay was employed to quantify disruption of CD28 binding to its native ligand, B7-1 (CD80), across a range of compound concentrations. In this assay, recombinant CD28 was immobilized on 96-well plates and incubated with biotinylated B7-1 in the presence of increasing concentrations of each compound. Residual ligand binding was detected using streptavidin–HRP and a chemiluminescent substrate. Dose–response curves revealed that both compounds effectively inhibited CD28:B7-1 interactions. Compound **5MS-5** revealed moderate inhibitory activity with an IC_50_ of 22.4 ± 6.8 μM (Figure S1A), while **19MS-5** showed stronger inhibition with an IC_50_ of 7.83 ± 4.80 μM (Figure S1B). These results confirm that biophysically validated CD28 binders can translate into functionally active inhibitors of T cell co-stimulation. Encouraged by these data, we advanced compounds **5MS-5** and **19MS-5** for further investigation in cellular models to assess their activity in more complex and physiologically relevant settings.

### Cellular CD28 Blockade Assay

To establish a direct link between CD28 binding and downstream functional activity, we assessed whether our top compounds (**5MS-5** and **19MS-5**) could inhibit CD28–B7 interactions in a cellular setting. We employed the CD28 Blockade Bioassay (Promega, Cat. #JA6101)—a luminescence-based reporter assay that captures T cell co-stimulatory signaling in response to CD28 engagement. In this system, CD28-expressing Jurkat T cells containing a TCR/CD3-CD28-responsive luciferase reporter are co-cultured with Raji antigen-presenting cells that endogenously express B7 ligands. The resulting luminescence reflects CD28-driven costimulation and is specifically reduced by compounds that block the CD28–B7 interaction. Remarkably, both compounds exhibited dose-dependent inhibition of CD28-mediated luminescence, consistent with their direct engagement of CD28 observed in previous binding assays. Compound **5MS-5** showed an IC_50_ of 10.32 ± 3.27 μM (Figure S2), while **19MS-5** exhibited the most potent activity with an IC_50_ of 5.20 ± 1.30 μM (Figure 6).

**Figure 6.**
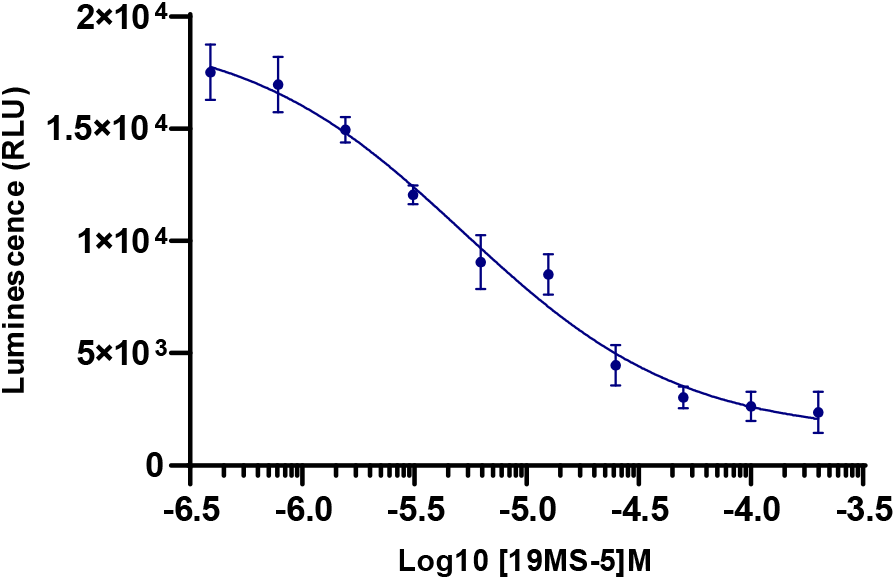
Functional inhibition of CD28–B7 signaling by 19MS-5 in a bioluminescent reporter assay. The CD28 Blockade Bioassay (Promega, Cat. #JA6101) was employed to assess the ability of **19MS-5** to inhibit CD28–B7 interactions in a co-culture system of CD28 Effector Cells (Jurkat) and aAPC/Raji Cells. The compound was evaluated in a 10-point dose–response format, and luminescence was quantified using the Bio-Glo™ Luciferase Assay System. Dose–response curve was fitted using nonlinear regression (four-parameter logistic model) in GraphPad Prism. Data are presented as mean ± SEM from n = 5 independent experiments.

### Computational Analysis of the Binding Mode at the CD28 Interface

To explore the structural basis of ligand engagement with CD28, we performed molecular docking of four compounds (**5MS, 19MS, 5MS-5**, and **19MS-5**) targeting a putative secondary binding pocket on the protein. Each ligand exhibited a distinct binding mode characterized by specific hydrogen bonding and hydrophobic interaction profiles (Figure 7). Compound **5MS** (Figure 7A–B) formed two hydrogen bonds with ASN107 and Ser110, stabilizing the ligand within the pocket. **19MS-5** (Figure 7C–D) demonstrated the most extensive interaction network, establishing four hydrogen bonds with Arg34, Cys94, Lys95, and Asn107, in addition to numerous hydrophobic contacts. **5MS-5** (Figure 7E–F) formed three hydrogen bonds involving Lys95, Asp106, and Asn107, while **19MS** (Figure 7G–H) engaged Asp106, Asn107, and Lys107 through hydrogen bonding, supported by favorable hydrophobic interactions.

**Figure 7.**
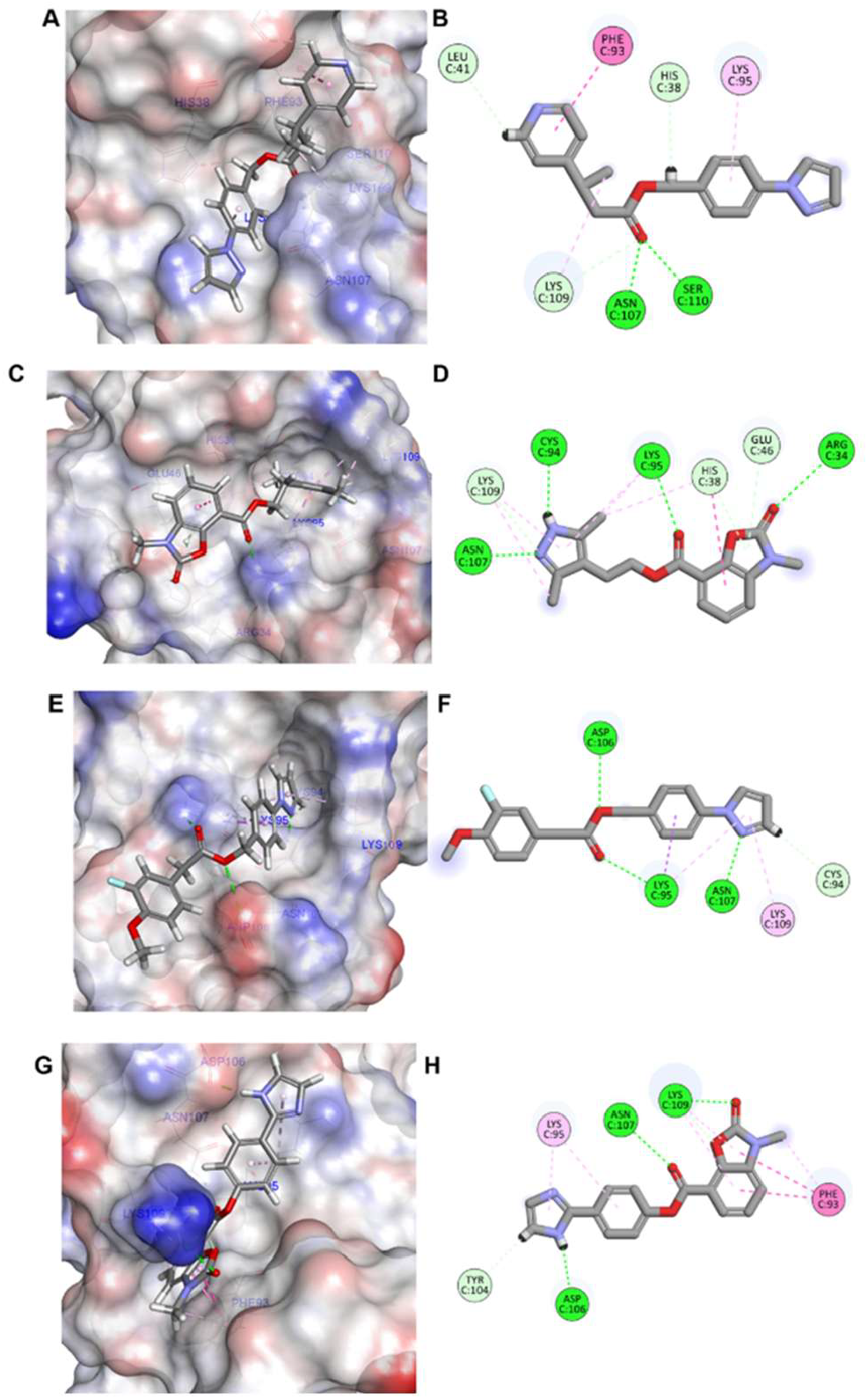
Molecular docking analysis of four compounds with CD28 protein using induced fit docking protocol. **(A-B)** Compound **5MS** docked into CD28, showing surface view **(A)** and detailed interaction diagram **(B). (C-D)** Compound **19MS-5** binding mode, displaying surface representation **(C)** and interaction network **(D). (E-F)** Compound **5MS-5** docked configuration, illustrated through surface view **(E)** and binding interactions **(F). (G-H)** Compound **19MS** binding analysis, showing surface representation **(G)** and detailed interactions **(H)**. In all panels, the protein surface is colored according to electrostatic potential (blue: positive, red: negative, white: neutral), and green dashed lines indicate hydrogen bonds or favorable electrostatic interactions. The ligands are represented as gray stick models, and interacting residues are labeled with their three-letter amino acid codes and residue numbers. Favorable interactions are color-coded as follows: green, hydrogen bonds; light green, carbon–hydrogen interactions; dark pink, π–π stacking interactions; purple, π–sigma interactions; light pink, hydrophobic interactions. All docking studies were performed using Maestro Schrödinger (version 2023.2) with induced fit protocols to account for protein flexibility during ligand binding.

These docking results underscore the importance of Asn107 as a common anchoring residue across all compounds, suggesting a shared binding mode and identifying it as a critical hot spot for ligand recognition. Furthermore, all four ligands contain a conserved pyrazole moiety, which appears to induce a localized conformational change in CD28. This rearrangement creates a shallow groove that accommodates ligand binding and may play a role in ligand-induced allosteric modulation. This effect was particularly pronounced in the **5MS-5** and **19MS-5** complexes, highlighting the pyrazole group as a key pharmacophoric feature for future optimization.

To validate the docking predictions and assess the dynamic stability of the ligand–CD28 complexes, we performed molecular dynamics (MD) simulations. Root-meansquare deviation (RMSD) analyses revealed striking differences in complex stability (Figure 8). The unbound CD28 protein (blue line) exhibited significant conformational fluctuations, especially after 25 ns, consistent with its inherent flexibility. In contrast, **19MS-5** (green line) and **19MS** (magenta line) maintained stable RMSD values consistently below 1.5 Å-, indicating robust binding and minimal conformational drift. These two compounds demonstrated the most stable protein-ligand complexes, which closely aligns with their functional potency observed in biological assays. Conversely, the CD28 complexes with **5MS** and **5MS-5** displayed greater RMSD fluctuations throughout the simulation period, suggesting weaker dynamic stability (Figure 8). This observation is in agreement with their reduced biological activity and higher binding K_d_ values, further supporting the reliability of the in-silico predictions. The correlation between MD stability, hydrogen bonding density, and experimental activity reinforces the value of structure-based modeling in guiding compound prioritization.

**Figure 8.**
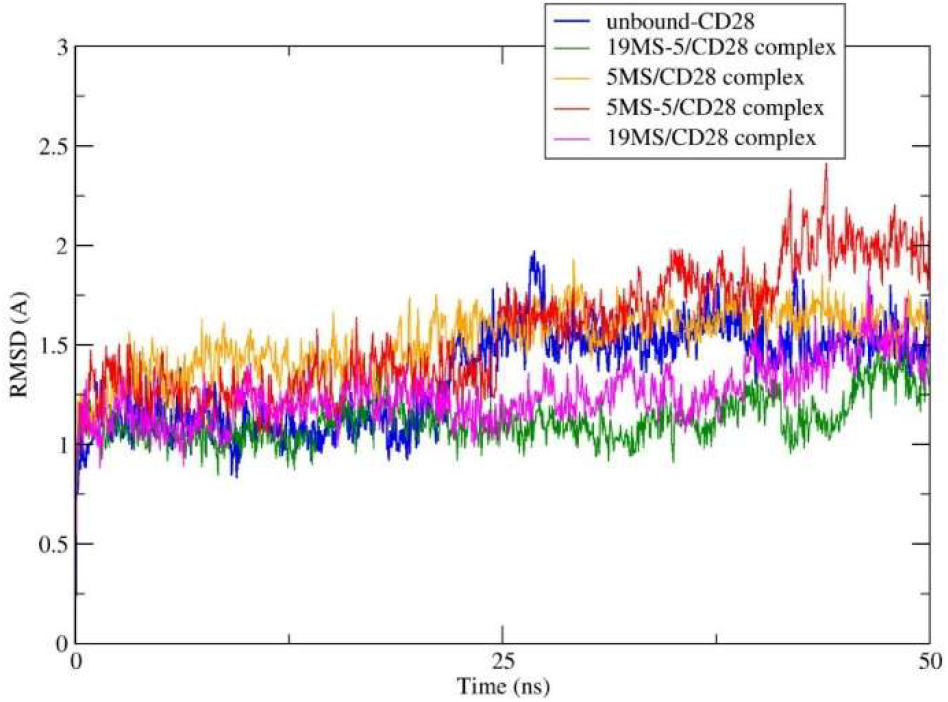
RMSD analysis of CD28 protein-ligand complexes during 50 ns molecular dynamics simulations.

Taken together, the integrated docking and MD simulation data highlight **19MS-5** as the most promising lead compound, exhibiting an optimal balance of hydrogen bonding, hydrophobic stabilization, conformational adaptation, and dynamic stability. These results provide valuable mechanistic insights into the molecular basis of CD28 modulation, establishing a solid foundation for the rational optimization of this scaffold aimed at enhancing ligand affinity and functional inhibition. By elucidating the interaction networks and conformational dynamics of these hits, this study offers a robust framework for advancing CD28-targeted immunomodulators.

### Preliminary PK evaluation of 5MS-5 and 19MS-5

To evaluate their drug-like properties, we performed a preliminary in vitro PK evaluation of **5MS-5** and **19MS-5** (Table 1). The LogD7.4 values of 3.31 for **5MS-5** and 1.87 for **19MS-5** indicate moderate lipophilicity, aligning well with their observed solubility and permeability. Both compounds displayed favorable aqueous solubility profiles, with kinetic solubility in PBS containing 1% DMSO measured at 97.4 μM for **5MS-5** and exceeding 100 μM for **19MS-5**. In biorelevant media (FaSSIF), **5MS-5** showed a solubility of 89.3 μM, while **19MS-5** exhibited slightly higher solubility at 124 μM, indicating good compatibility with physiological gastrointestinal conditions and potential for oral administration. Passive membrane permeability, assessed via PAMPA, revealed *P*_*app*_ values of 1.5 × 10^−6^ cm/s for **5MS-5** and 2.1 × 10^−6^ cm/s for **19MS-5**, both within the range expected for orally bioavailable compounds. Stability studies in simulated gastric fluid (pH 1.2) and intestinal fluid (pH 6.8) showed that both compounds were chemically robust, maintaining >90% (**5MS-5**) and >85% (**19MS-5**) integrity after 2 hours of incubation in each environment.

**Table 1.** In vitro PK profile of 5MS-5 and 19MS-5.

| PK parameter | 5MS-5 | 19MS-5 |
| --- | --- | --- |
| LogD<sub>7.4</sub><sup>a</sup> | 3.31 | 1.87 |
| Kinetic Solubility<br>1% DMSO/PBS<br>[μM]<sup>b</sup> | 97.4 | >100 |
| Solubility in FaSSIF<br>[μM] | 89.3 | 124 |
| PAMPA *P*<sub>app</sub> [cm/s] | 1.5 × 10<sup>-6</sup> | 2.1 × 10<sup>-6</sup> |
| Stability in simulated gastric fluid<br>(pH 1.2, 2h) | >90% | >85% |
| Stability in simulated intestinal fluid<br>(pH 6.8, 2h) | >90% | >85% |
| *t*<sub>1/2</sub> [min] mouse liver microsomes / CL<sub>int</sub><sup>d</sup> | 92.5 ± 4.1 / 25.1 ± 2.8 | 97.1 ± 3.7 / 18.3 ± 1.6 |
| *t*<sub>1/2</sub> [min] human liver microsomes<sup>c</sup> / CL<sub>int</sub><sup>d</sup> | 101.3 ± 8.2 / 16.4 ± 0.9 | 110.2 ± 5.4 / 14.9 ± 1.2 |
| *t*<sub>1/2</sub> [min] mouse plasma | >150 | >150 |
| *t*<sub>1/2</sub> [min] human plasma | >150 | >150 |
| *t*<sub>1/2</sub> [min] PBS | >300 | >300 |
| PPB human [%]<sup>c</sup> | 98.5 ± 0.47 | 91.1 ± 0.82 |
| IC<sub>50</sub> against WI-38 [μM] | >100 | >100 |
| IC<sub>50</sub> against HS 27 [μM] | >100 | >100 |
| IC<sub>50</sub> against hERG [μM] | >50 | >50 |
| CYP3A4 inhibition (% at 10 μM) | 13.4 | 14.1 |
| CYP2D6 inhibition (% at 10 μM) | 4.3 | 5.2 |
| CYP2C9 inhibition (% at 10 μM) | 11.2 | 10.1 |
| CYP2C19 inhibition (% at 10 μM) | 7.7 | 6.3 |

<sup>a</sup>LogD (pH = 7.4) the distribution coefficient at physiological pH 7.4. <sup>b</sup>Kinetic solubility measured in 1% DMSO/PBS (pH 7.4) by UV–vis spectrophotometry (n=3). <sup>c</sup>Values are mean ± SD (n=3). <sup>d</sup>Intrinsic clearance expressed in μL/(min × mg) protein. Values are mean ± SD (n=3).

Microsomal stability studies demonstrated moderate intrinsic clearance in both species. In mouse liver microsomes, **5MS-5** had a half-life of 92.5 ± 4.1 min (CL_int_ = 25.1 ± 2.8 μL/min/mg), while **19MS-5** exhibited a slightly longer half-life of 97.1 ± 3.7 min (CL_int_ = 18.3 ± 1.6 μL/min/mg). Similar trends were observed in human liver microsomes, where **5MS-5** and **19MS-5** achieved half-lives of 101.3 ± 8.2 min and 110.2 ± 5.4 min, respectively. These data suggest both compounds are metabolically stable and not rapidly cleared via hepatic metabolism. Both compounds showed excellent plasma stability in human and mouse plasma (t_1_/_2_ > 150 min) and remained chemically stable in PBS for over 5 hours (t_1_/_2_ > 300 min). Plasma protein binding (PPB) was high for **5MS-5** (98.5 ± 0.47%) and moderately high for **19MS-5** (91.1 ± 0.82%), indicating strong association with serum proteins that may affect free drug availability but could also prolong in vivo half-life.

In safety liability profiling, neither compound showed cytotoxicity against normal human cell lines WI-38 and HS-27 up to 100 μM. Additionally, both **5MS-5** and **19MS-5** showed weak inhibition of the hERG potassium channel (IC_50_ > 50 μM), suggesting a low risk of cardiotoxicity. Evaluation against major cytochrome P450 isoforms revealed minimal inhibition at 10 μM for CYP3A4 (13.4% and 14.1%, respectively), CYP2D6 (4.3% and 5.2%), CYP2C9 (11.2% and 10.1%), and CYP2C19 (7.7% and 6.3%), indicating low potential for drug–drug interactions. Together, these in vitro data highlight the favorable PK, physicochemical, and safety profiles of **5MS-5** and **19MS-5**. Their combination of metabolic stability, solubility, permeability, and low off-target liability supports their advancement as lead CD28-targeted small molecules for further optimization and in vivo evaluation.

### Evaluation of T Cell Activation in a Tumor–PBMC Coculture System

To investigate the immunomodulatory effects of our lead CD28 inhibitors in a physiologically relevant context, we utilized a 3D co-culture model comprising IFN-γ–primed A549 tumor spheroids and human PBMCs. Co-cultures were stimulated with suboptimal levels of anti-CD3 to enable co-stimulation-dependent T cell activation driven by tumor-expressed CD80/CD86. Under these conditions, robust T cell activation (Figure 9) was observed, evidenced by elevated secretion of canonical markers such as IFN-γ, IL-2, and soluble CD69 (sCD69). Treatment with **5MS-5** and **19MS-5** resulted in concentration-dependent suppression of T cell activation, indicating effective inhibition of CD28-mediated co-stimulation (Figure 9). Notably, **19MS-5** exhibited stronger activity across all readouts compared to **5MS-5**, consistent with its superior biochemical potency. We used the clinical-stage CD28 antagonist FR104 as a positive control in this study. These results validate both compounds as inhibitors of CD28-driven T cell responses in a human tumor immune microenvironment model, with **19MS-5** emerging as the more promising candidate for downstream optimization.

**Figure 9.**
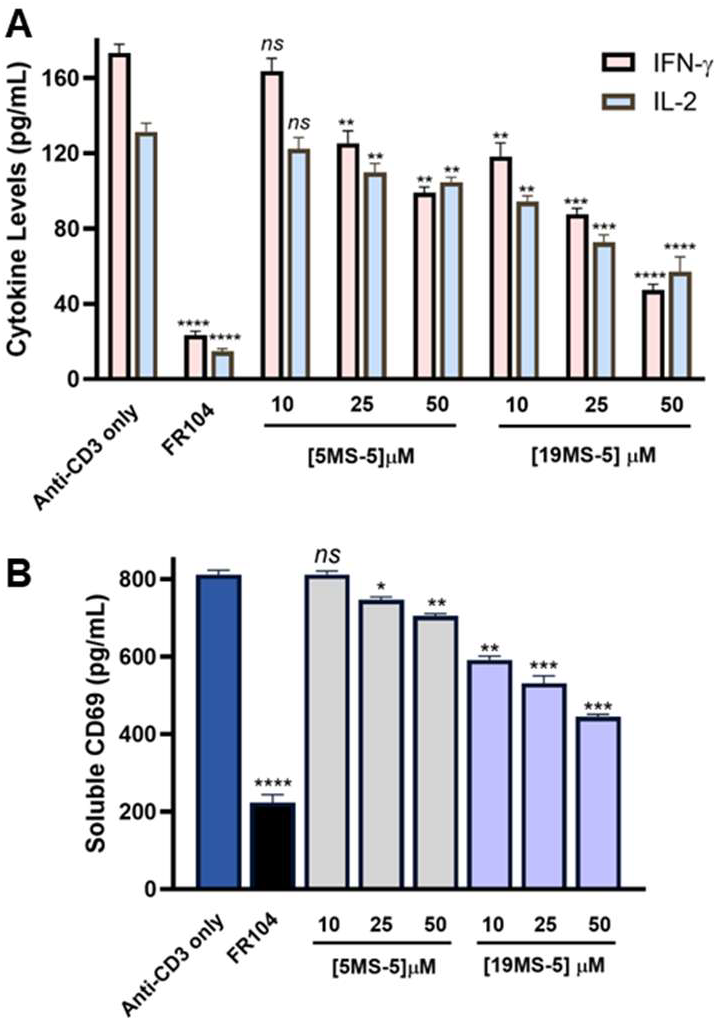
Dose-dependent inhibition of T cell activation markers in tumor–PBMC co-culture. **(A)** Soluble IFN-γ and IL-2 levels were quantified after 48-hour co-culture of A549 tumor spheroids with human PBMCs (E:T ratio 5:1) in the presence of anti-CD3 (0.3 μg/mL) and test compounds: FR104 (10 μg/mL), **5MS-5** (10, 25, 50 μM), or **19MS-5** (10, 25, 50 μM). **(B)** Soluble CD69 levels measured by ELISA after 48-hour co-culture of A549 tumor spheroids with human PBMCs (E:T ratio 5:1) in the presence of anti-CD3 (0.3 μg/mL) and test compounds: FR104 (10 μg/mL), **5MS-5** (10, 25, 50 μM), or **19MS-5** (10, 25, 50 μM). Data represent mean ± SD of n = 3 independent wells. Statistical comparisons to the “Anti-CD3 only” group were performed using one-way ANOVA followed by Dunnett’s post-hoc test. ns denotes non-significant, * *p* < 0.05, ** *p* < 0.01, *** *p* < 0.001, and **** *p* < 0.0001 relative to anti-CD3.

### Human PBMC–Mucosal Co-culture Model

To further validate the functional activity of **19MS-5**, we employed a human co-culture system that models immune-epithelial interactions at mucosal surfaces. In this assay (Figure 10A), fully differentiated airway epithelial tissues (MucilAir™, Epithelix) were overlaid with primary human PBMCs and stimulated with a combination of anti-CD3 and soluble anti-CD28 antibodies to drive T cell co-stimulation. This model recapitulates key features of the mucosal immune microenvironment, including epithelial barrier integrity, local cytokine gradients, and immune–epithelial crosstalk. Cytokine levels were quantified in the apical compartment after 48 hours of co-culture to assess the impact of pharmacological CD28 blockade.

**Figure 10.**
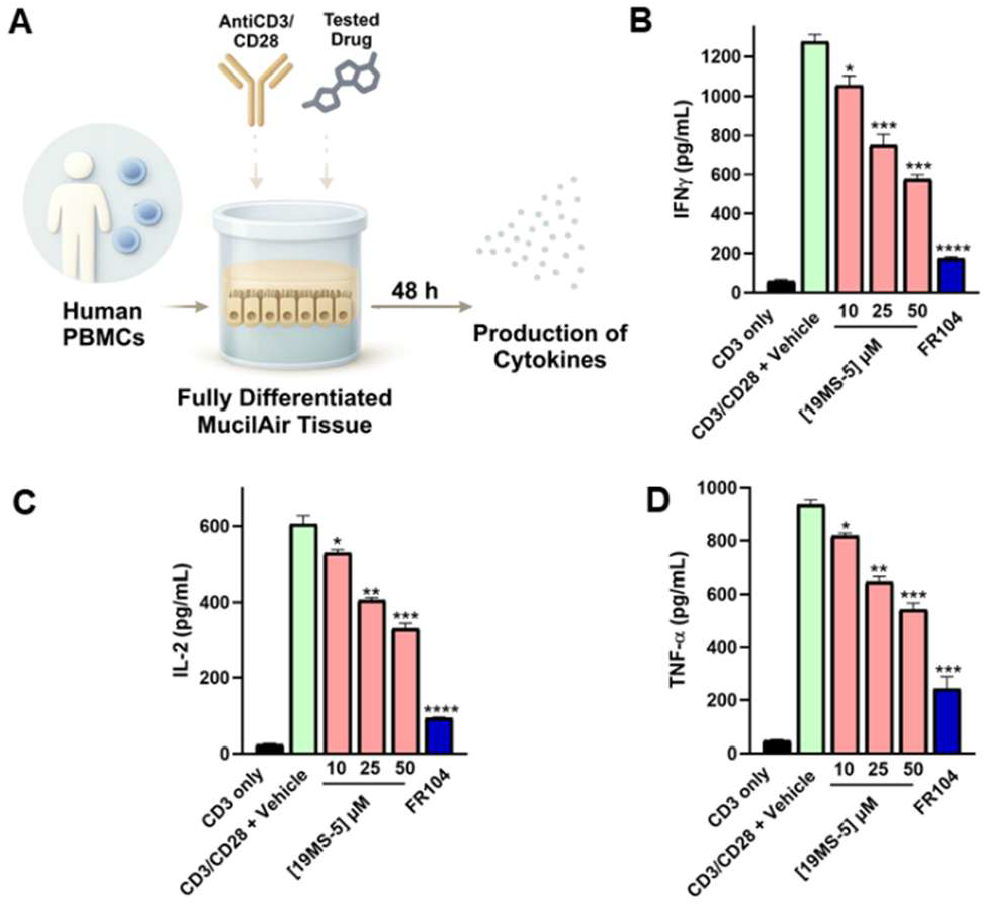
19MS-5 suppresses CD28-dependent cyto-kine release in a human PBMC–mucosal co-culture model. **(A)** Schematic overview of the experimental setup. Human PBMCs were overlaid onto fully differentiated Mu-cilAir™ tissue (Epithelix) and stimulated with plate-bound anti-CD3 and soluble anti-CD28 for 48 h in the presence of vehicle (0.1% DMSO), **19MS-5** (10, 25, or 50 μM), or FR104 (10 μg/mL). **(B–D)** Quantification of IFN-γ, IL-2, and TNF-α secretion (pg/mL) in apical supernatants by ELISA. **19MS-5** suppressed CD28-induced cytokine production in a dose-dependent manner, with levels at 50 μM comparable to FR104. Statistical comparisons to the “CD3/CD28 + Vehicle” group were performed using one-way ANOVA followed by Dunnett’s post-hoc test. * *p* < 0.05, ** *p* < 0.01, *** *p* < 0.001, and **** *p* < 0.0001 relative to “CD3/CD28 + Vehicle” group.

Stimulation with anti-CD3/CD28 led to robust secretion of proinflammatory cytokines, including IFN-γ, IL-2, and TNF-α, indicative of strong CD28-mediated T cell activation. Treatment with **19MS-5** resulted in dose-dependent suppression of all three cytokines. At lower concentrations, a modest reduction was observed, while higher concentrations of **19MS-5** substantially decreased cytokine output (Figures 10B-C). As a benchmark, we included FR104, a clinical-stage CD28 antagonist, to validate assay performance. These findings demonstrate that **19MS-5** maintains potent functional activity in a primary human tissue model, mirroring its effects in tumor–PBMC co-culture assays. The consistency of response across distinct co-culture systems underscores the ability of **19MS-5** to effectively inhibit CD28-dependent immune activation in physiologically relevant settings.

## CONCLUSIONS

In this study, we report the discovery and characterization of first-in-class small molecule allosteric antagonists targeting the CD28 co-stimulatory receptor, a key regulator of T cell activation with limited chemical tractability to date. Using a SEC-coupled AS-MS platform, we enabled label-free, solution-phase identification of CD28-binding compounds from a chemical library with structural diversity for Enamine. This was followed by TRIC-based validation and SAR-by-catalog optimization, yielding two lead compounds (**5MS-5** and **19MS-5**) that directly bind CD28 and disrupt its interaction with B7 ligands. Both compounds demonstrated low micromolar affinities, favorable pharmacokinetic properties (including high solubility, metabolic stability, and membrane permeability), and minimal off-target liabilities. Functionally, **5MS-5** and **19MS-5** potently suppressed cytokine production in human tumor–PBMC and mucosal–PBMC co-culture models, validating their inhibitory activity in physiologically relevant settings. Collectively, our findings establish a chemical starting point for the development of orally bioavailable CD28-targeted immunotherapies and underscore the power of SEC-coupled AS-MS and TRIC-based platforms for ligand discovery against challenging immunomodulatory targets. This work opens new avenues in immunopharmacology beyond antibodies and fusion proteins. Our findings not only provide immediate chemical tools to interrogate CD28 signaling but also lay the groundwork for developing small molecule immunotherapies with tunable potency, enhanced selectivity, and reduced immunogenicity.

## EXPERIMENTAL

### SEC-ASMS Screening and LC-MS Analysis

Recombinant CD28 protein was obtained from Acro Biosystems (Cat # CD8-H52H3) and used for all affinity selection experiments. The construct comprised the extracellular domain of human CD28 (cross-reactive with cynomolgus and rhesus macaque), fused to a C-terminal His tag. Dimeric assembly of the protein was confirmed via multi-angle light scattering (MALS). The protein was supplied at a concentration of 0.6 mg/mL (15 μM, molecular weight ~40 kDa) in PBS (pH 7.4) and was stored at −80°C until use. For all experiments, the protein was freshly thawed and buffer-exchanged into ultrapure water for storage or 50 mM KH_2_PO4, pH 7.4, with 100 mM NaCl for incubation and screening steps.

The compound library consisted of 6,336 chemically diverse small molecules, screened initially in pools of 1,056 compounds per mixture. For the primary SEC-ASMS screen, each pool was tested at a final concentration of 1 μM per compound, with 2.5 μM CD28 protein. Compounds were incubated with the protein in 50 mM KH_2_PO4 buffer (pH 7.4) containing 100 mM NaCl. All incubations were performed at 4°C with minimal agitation to preserve weak or transient interactions. For hit confirmation (retest) screens, candidate pools were reassembled into focused mixes of ~10 compounds each and tested at an elevated CD28 concentration of 5 μM while maintaining 1 μM per compound. This format allowed for higher resolution detection of reproducible binders and mitigation of competitive suppression within complex pools.

AS-MS was carried out using a two-dimensional SEC-LC– MS approach. After compound incubation, protein–ligand mixtures were directly injected onto a PolyLC size exclusion column (2.1 × 50 mm, 5 μm) operating at 8°C with PBS or 50 mM KH_2_PO4, pH 7.4, as the mobile phase at a flow rate of 1.0. mL/min. Protein–ligand complexes were separated based on molecular weight, and the high-molecular-weight fractions were collected online and passed to the second-dimension reversed-phase LC-MS. For second-dimension separation, samples were loaded onto an Agilent RRHD C18 column (2.1 × 50 mm, 1.7 μm) maintained at 60°C. The mobile phases were 0.2% formic acid in water with 50 mM ammonium acetate (aqueous) and 0.2% formic acid in acetonitrile (organic). The gradient was set as follows: isocratic hold at 2% B from 0–0.5 min, linear ramp to 95% B from 0.5–2.5 min, hold at 95% B until 3.5 min, and return to 2% B by 3.6 min, followed by equilibration until 5.0 min. The flow rate was 0.3 mL/min, and the injection volume was 40 μL. This two-step separation ensured robust desalting, peak resolution, and compatibility with mass spectrometry.

### Hit Identification Criteria

MS data were collected in positive ion mode using a high-resolution time-of-flight mass spectrometer calibrated for <5 ppm mass accuracy. Each compound detected in the protein-incubated (P^+^) samples was compared to the corresponding P^−^ (no protein) control for evidence of enrichment. Hit selection criteria included: (1) accurate monoisotopic mass within ±5 ppm of theoretical value, (2) consistent retention time across duplicate P^+^ runs, (3) reproducible isotopic envelope, and (4) significantly reduced or absent signal in the P^−^ sample. Signal intensities were averaged across replicates to ensure robustness. In summary, in the primary screen, 52 compounds were identified as CD28 binders from a total of 6,336 compounds, resulting in a primary hit rate of 0.82%. These hits were then retested in smaller pools under higher protein concentration, leading to confirmation of 25 ligands, corresponding to a validated hit rate of 0.39%. The screening and validation strategy demonstrated high specificity and reproducibility, providing a solid foundation for downstream biophysical and functional characterization.

### Dianthus TRIC Assay

To enable high-throughput identification of small molecules that bind human CD28, we employed the TRIC (temperature-related intensity change) assay using the Dianthus NT.23 Pico instrument (NanoTemper Technologies). His-tagged recombinant human CD28 protein (Acro Biosystems) was fluorescently labeled with RED-tris-NTA 2nd Generation dye (NanoTemper, Cat. #MO-L018) following the manufacturer’s protocol. The labeling mixture consisted of 40 nM CD28 and 20 nM dye in assay buffer composed of phosphate-buffered saline (PBS), pH 7.4, supplemented with 0.005% Tween-20. The solution was incubated in the dark for 30 minutes at ambient temperature.

For single-point screening, labeled CD28 was mixed in a 1:1 volume ratio with individual test compounds (final compound concentration: 100 μM in 2% DMSO) in a 384-well plate format. Following a 15-minute incubation at room temperature in the dark, samples were centrifuged briefly (1 min at 1000×g), and 20 μL of each mixture was loaded into the instrument. Fluorescence emission at 670 nm was monitored for 1 second before and 5 seconds after IR laser activation. The normalized TRIC signal (Fnorm) was calculated as the ratio of post-laser (Fhot) to pre-laser (Fcold) fluorescence intensities. Human CD80 (2 μM) served as a positive control for CD28 binding, while 2% DMSO was used as a negative control. Each test compound was analyzed in triplicate, and controls were placed at regular intervals across the screening plate to ensure assay consistency.

### Quantitative Binding Affinity Determination Using Monolith

To quantitatively characterize binding affinities of selected hit compounds, microscale thermophoresis (MST) measurements were performed using the Monolith NT.115 platform (NanoTemper Technologies). His-tagged CD28 protein (Acro Biosystems) was labeled with RED-tris-NTA 2nd Generation dye using the Monolith His-Tag Labeling Kit (Cat. #MO-L018) according to the manufacturer’s instructions. A mixture containing 100 nM CD28 and 50 nM dye in PBS buffer (pH 7.4, 0.005% Tween-20) was incubated for 30 minutes at room temperature in the dark.

Binding titrations were carried out using 12-point serial dilutions of each test compound, starting from 500 μM in 2% DMSO, to prepare 2× stocks. These were mixed in a 1:1 ratio with the labeled CD28 solution to yield a final reaction volume of 15 μL per sample. After a 15-minute incubation at room temperature in the dark, samples were loaded into Monolith capillaries (Cat. #MO-K022) and analyzed at 25 °C using 40% LED power and medium MST power settings. Fnorm values were calculated as the fluorescence intensity ratio after vs. before IR laser heating. All compounds were tested in four technical replicates. Binding affinities (Kd values) were derived from three independent experiments using MO. Affinity Analysis software and GraphPad Prism 10, based on dose–response fitting with standard binding models.

### Assessment of CD28–CD80 Interaction Inhibition via Competitive ELISA

To determine the functional ability of small molecules to inhibit the interaction between CD28 and its natural ligand CD80 (B7-1), we performed a competitive ELISA using the CD28:B7-1 [Biotinylated] Inhibitor Screening Assay Kit (BPS Bioscience, Cat. #72007), following the manufacturer’s protocol. Briefly, 96-well plates were coated with recombinant human CD28 protein (2 μg/mL in PBS, 50 μL per well) and incubated overnight at 4 °C. The following day, wells were washed with 1× Immuno Buffer and blocked with the supplied Blocking Buffer for 1 hour at room temperature. Test compounds were prepared at varying concentrations and incubated with biotinylated CD80 (5 ng/μL final concentration) in Blocking Buffer for 1 hour at room temperature to enable competitive binding to the immobilized CD28. Control wells without CD28 coating were included to assess background signal from the biotinylated ligand (ligand control), while wells treated with inhibitor buffer in place of compound served as negative controls. After the competition step, plates were washed thoroughly and incubated with Streptavidin-HRP (1:1000 dilution in Blocking Buffer) for 1 hour at room temperature. Following the final wash step, chemiluminescent substrate was added to each well, and luminescence was immediately measured using a multimode plate reader operating in luminescence detection mode. Inhibition curves were generated by plotting luminescence intensity against compound concentration, with each assay conducted in at least two independent experiments.

### SAR-by-Catalog and Affinity Validation of Analog Compounds

To explore SAR and improve the potency of initial CD28-binding compounds, we conducted an analog-based screening campaign using catalog-available derivatives sourced from Enamine. A total of 20 structurally related analogs were selected based on scaffold similarity to initial hits using substructure search and medicinal chemistry filters.

### CD28 Blockade Bioassay for Functional Inhibition and Serum Tolerance Assessment

A luminescence-based CD28 Blockade Bioassay (Promega, Cat. #JA6101) was adapted to a 96-well format to evaluate the ability of small molecules to inhibit CD28-mediated costimulatory signaling.^41^ Test compounds were prepared in a 10-point, 1:1 serial dilution series starting from 200 μM in assay buffer containing 1% DMSO. Each concentration (50 μL per well) was transferred into white 96-well assay plates. CD28 Effector Cells (Jurkat cells expressing a CD28- and TCR-responsive luciferase reporter) were seeded at 2 × 104 cells per well and preincubated with the diluted compounds or a positive control anti-CD28 antibody (Promega, Cat. #K1231) for 5 minutes at room temperature. Subsequently, artificial antigen-presenting cells (aAPC/Raji cells), which endogenously express CD80/CD86, were added at 2 × 104 cells per well. The plates were incubated for 5 hours at 37 °C in a humidified 5% CO_2_ incubator. After incubation, 50 μL of Bio-Glo™ Luciferase Assay Reagent (Promega) was added to each well, and luminescence was measured using a GloMax® Discover System (Promega). Dose–response curves were generated from normalized luminescence values and analyzed using four-parameter logistic regression in GraphPad Prism 10 to determine percent inhibition and IC_50_ values. To assess compatibility with physiological conditions, the assay was additionally performed in the presence of pooled human serum (up to 10%). No significant interference was observed, demonstrating the robustness and matrix tolerance of the assay under near-physiological conditions.

### In Silico Methods for Binding Mode and Stability Analysis

Molecular docking studies were conducted using Maestro Schrödinger software (version 2023.2) to evaluate the binding interactions of four compounds (**5MS, 5MS-5, 19MS** and **19MS-5**) with the CD28 protein. The target compounds were prepared using the LigPrep module, which optimized their three-dimensional structures, assigned appropriate ionization states, and generated conformers suitable for docking calculations. The CD28 protein structure was processed through the Protein Preparation Wizard module, which involved hydrogen addition, optimization of side chain conformations, and energy minimization to ensure structural integrity. The induced fit docking protocol was employed to account for protein flexibility during ligand binding, allowing both the ligand and receptor to adapt their conformations for optimal complementarity. The compounds were specifically docked into the secondary binding pocket of CD28, which had been previously characterized^16^ as possessing favorable physicochemical properties for small molecule targeting and drug development.

Five 50ns MD simulations were performed using the Desmond simulation package to evaluate the stability and dynamic behavior of the CD28 protein-ligand complexes of the identified hits in comparison to the unbound CD28 protein. The MD simulations followed the established protocol described in our previous study, ensuring consistency and reproducibility of the computational approach. All four compound-CD28 complexes (**5MS, 5MS-5, 19MS** and **19MS-5**) along with the unbound CD28 protein were subjected to MD simulations under identical conditions. RMSD analysis was performed to monitor structural changes and binding stability throughout the simulation trajectories. The docking results and MD trajectories were visualized using Discovery Studio Visualizer (version 24.1.0), while RMSD plots were generated and analyzed using Grace software to quantify the structural fluctuations of each system over time.

### HPLC Purity

The purity of **5MS, 5MS-5, 19MS** and **19MS-5** was confirmed to be >95% using HPLC (Figure S3-6). HPLC analysis was performed with a reversed-phase column Phenomenex Gemini, C18 (250 mm × 4.60 mm, 5 μm) on an HPLC Agilent system. The mobile phase used was an acetonitrile-H_2_O gradient and a 1 mL/min flow rate. UV absorption at 210 and 254 nm was used to monitor the method.

### PK Studies

The preliminary evaluation of PK parameters for **5MS-5** and **19MS-5** was performed as previously reported by us.^38^

### Evaluation of T Cell Activation in Tumor-PBMC Coculture

To assess the dose-dependent impact of CD28 inhibition on T cell activation in a translational human system, we employed a 3D tumor–PBMC co-culture assay. Tumor spheroids were generated by seeding A549 (or MDA-MB-231) cells in ultra-low attachment 96-well plates. After 48 hours, spheroids were treated with recombinant human interferon-gamma (IFN-γ, 50 ng/mL) for 24 hours. Freshly thawed human peripheral blood mononuclear cells (PBMCs) were added at a 5:1 effector-to-target (E:T) ratio, and co-cultures were stimulated with soluble anti-CD3 antibody (0.3 μg/mL, clone OKT3) to model TCR activation. Compounds were added at the initiation of co-culture. FR104 (10 μg/mL), a reference anti-CD28 Fab’ biologic, served as a positive control. The test compounds **5MS-5** and **19MS-5** were evaluated at three concentrations (10, 25, and 50 μM). Vehicle and anti-CD3–only conditions were included as controls. After 48 hours, supernatants were harvested and analyzed for IFN-γ and IL-2 levels using human-specific ELISA kits (BioLegend). Soluble CD69 (sCD69), released from activated T cells, was quantified using a human CD69 ELISA kit (ThermoFisher).

### CD28-Dependent Cytokine Release in a Human PBMC–Mucosal Co-culture Model

To evaluate the immunomodulatory activity of **19MS-5** in a mucosal-relevant human immune interface, we employed a commercially available human airway epithelial tissue model (MucilAir™, Epithelix) co-cultured with PBMCs. Cryopreserved human PBMCs (StemCell Technologies) were thawed and rested for 4 hours in RPMI-1640 medium supplemented with 10% heat-inactivated human AB serum, 2 mM L-glutamine, and 1% penicillin–streptomycin. MucilAir™ inserts were equilibrated in the manufacturer’s proprietary maintenance medium for 24 hours at 37 °C and 5% CO2. PBMCs were resuspended at 2 × 106 cells/mL and overlaid (200 μL per insert) onto the apical surface of the MucilAir™ tissue in 24-well Transwell plates. Co-cultures were stimulated with platebound anti-CD3 (1 μg/mL) and soluble anti-CD28 (1 μg/mL) in the presence of: vehicle control (0.1% DMSO), **19MS-5** (10, 25, or 50 μM), or FR104 (10 μg/mL; anti-CD28 Fab’ fragment). After 48 hours, apical supernatants were harvested and analyzed for IFN-γ, IL-2, and TNF-α concentrations using human ELISA kits (BioLegend) according to manufacturer instructions. Cytokine concentrations were interpolated from standard curves and expressed in pg/mL. All conditions were performed in triplicate (three independent experiments).

## ASSOCIATED CONTENT

### Supporting Information

The Supporting Information is available free of charge on the ACS Publications website.

Chemical structures of primary hit compounds, ELISA-based inhibition of CD28 interaction, CD28 cellular blockade assay, and HPLC traces (PDF)

## AUTHOR INFORMATION

### Author Contributions

The manuscript was written through contributions of all authors. All authors have given approval to the final version of the manuscript.

## ABBREVIATIONS

AS-MS: affinity selection–mass spectrometry
CD: cluster of differentiation
CL_int_: intrinsic clearance
DEL: DNA-encoded library
FaSSIF: fasted-state simulated intestinal fluid
IC_50_: halfmaximal inhibitory concentration
IFN-γ: interferon gamma
IL-2: interleukin-2
K_d_: equilibrium dissociation constant
LogD: logarithm of distribution coefficient
MD: molecular dynamics
MST: microscale thermophoresis
PBMC: peripheral blood mononuclear cell
PBS: phosphate-buffered saline
PK: pharma-cokinetics
PPB: plasma protein binding
sCD69: soluble cluster of differentiation 69
SAR: structure–activity relationship
SEC: size-exclusion chromatography
TCR: T cell receptor
TRIC: temperature-related intensity change.

## Notes

The authors declare no competing financial interests.

## ACKNOWLEDGMENTS

This work was supported by the National Institute of Diabetes and Digestive and Kidney Diseases (NIDDK) under grant number R01DK137299 (PI: Gabr).

