## Supporting Information for "Discovery of CD28-Targeted Small Molecule Inhibitors of T Cell Co-stimulation Using Affinity Selection-Mass Spectrometry (AS-MS) and Ex Vivo Validation"

*Electronic Supplementary Information*

| **Contents** |  |
| --- | --- |
| Chemical structures of the preliminary hit compounds identified by SEC-coupled AS-MS screening | S2 |
| Inhibition of CD28:B7-1 binding by 5MS-5 (A) and 19MS-5 (B) as measured by an ELISA-based CD28:B7-1 inhibitor screening assay | S7 |
| Functional inhibition of CD28–B7 signaling by 5MS-5 in a bioluminescent reporter assay | S7 |
| HPLC trace of compound **5MS** | S8 |
| HPLC trace of compound **5MS-5** | S8 |
| HPLC trace of compound **19MS** | S9 |
| HPLC trace of compound **19MS-5** | S9 |

**Table S1**. Chemical structures of the preliminary hit compounds identified by SEC-coupled AS-MS screening.

| **Compound name** | **Enamine Code** | **Chemical structure** |
| --- | --- | --- |
| **1MS** | Z1230942077 | 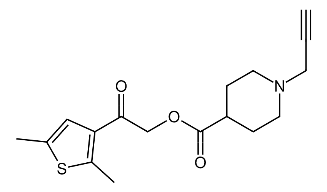 |
| **2MS** | Z1516160734 | 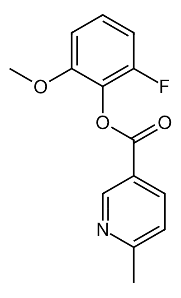 |
| **3MS** | Z740999256 | 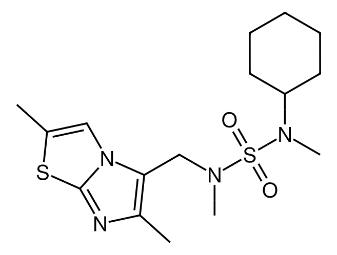 |
| **4MS** | Z1469064952 | 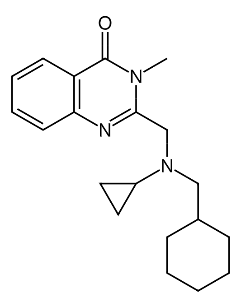 |
| **5MS** | Z1412833783 | 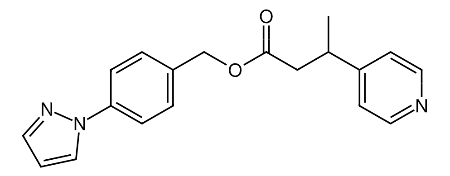 |
| **6MS** | Z1439189874 | 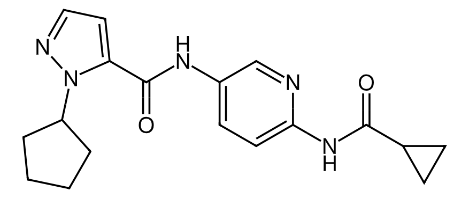 |
| **7MS** | Z971087312 | 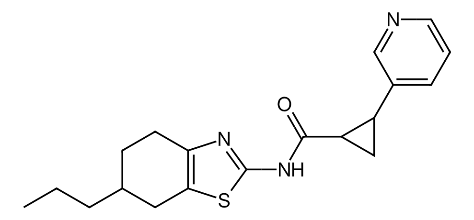 |
| **8MS** | Z1558830925 | 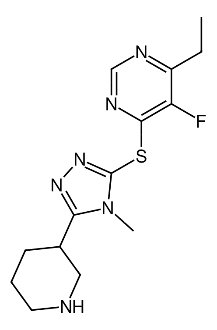 |
| **9MS** | Z1101149011 | 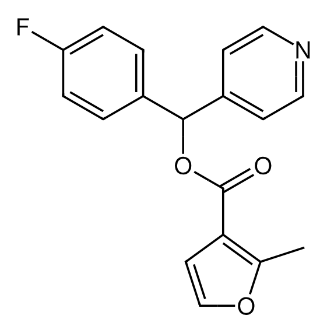 |
| **10MS** | Z1185860972 | 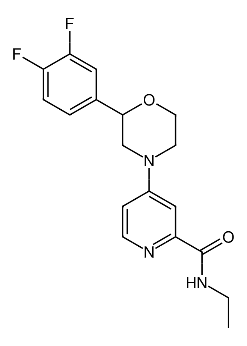 |
| **11MS** | Z1621796400 | 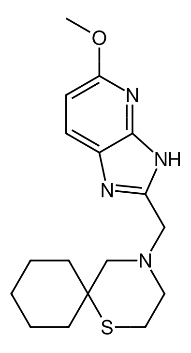 |
| **12MS** | Z1798321314 | 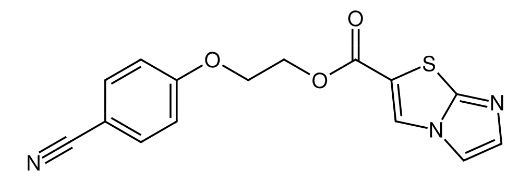 |
| **13MS** | Z1591059457 | 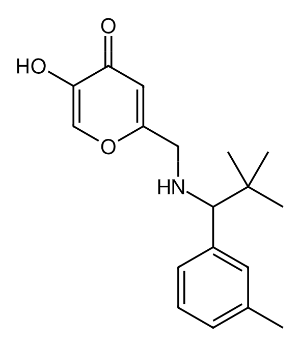 |
| **14MS** | Z1341536352 | 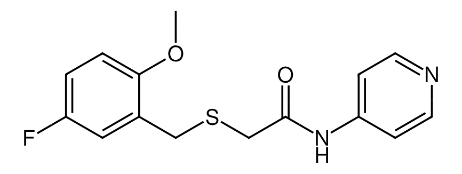 |
| **15MS** | Z1329374478 | 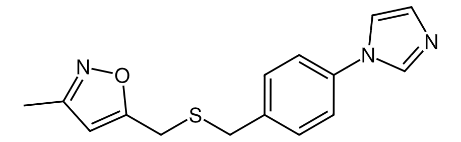 |
| **16MS** | Z1502794102 | 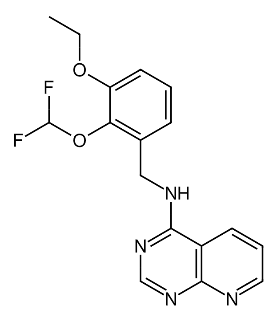 |
| **17MS** | Z55180110 | 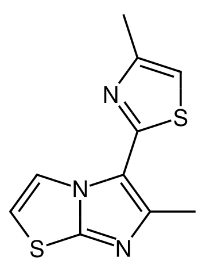 |
| **18MS** | Z237485898 | 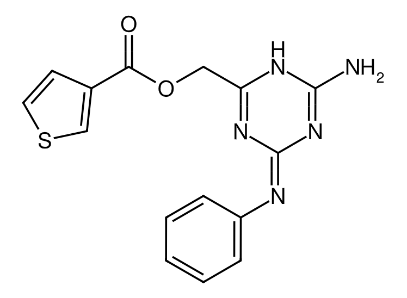 |
| **19MS** | Z1522726483 | 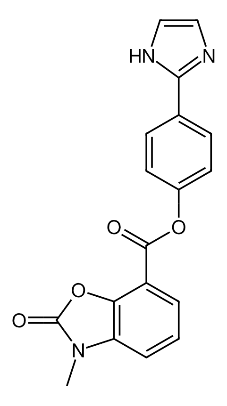 |
| **20MS** | Z1642773713 | 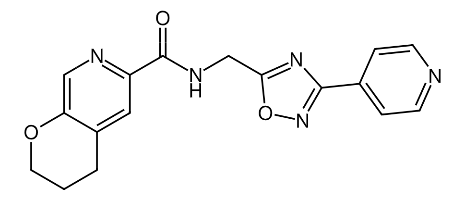 |
| **21MS** | Z450685474 | 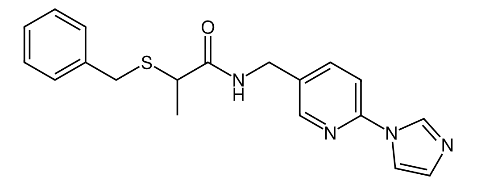 |
| **22MS** | Z1898856141 | 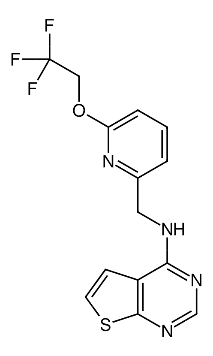 |
| **23MS** | Z1402651285 | 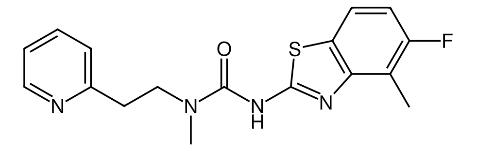 |
| **24MS** | Z1348439558 | 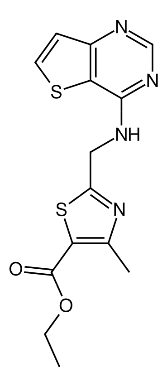 |
| **25MS** | Z1751965519 | 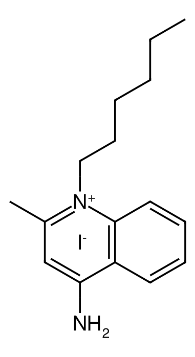 |

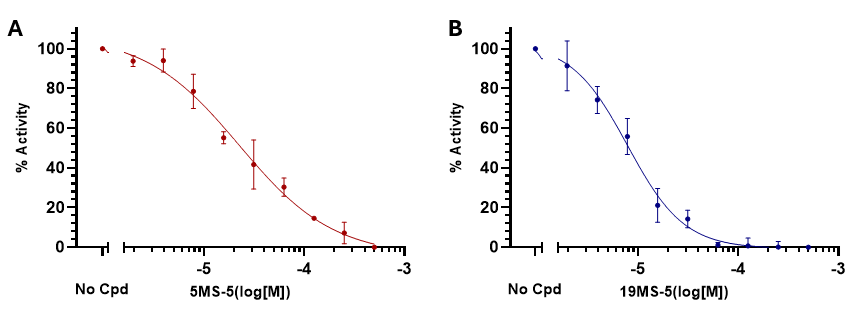

**Figure S1. Inhibition of CD28:B7-1 binding by 5MS-5 (A) and 19MS-5 (B) as measured by an ELISA-based CD28:B7-1 inhibitor screening assay. (A)** The structural analog **5MS-5** exhibited an IC_50_ of 22.4 ± 6.8 μM. **(B)** The analog **19MS-5** displayed an IC_50_ of 7.83 ± 4.8 μM. Data are presented as mean ± standard error mean (SEM) from n = 5 independent experiments.

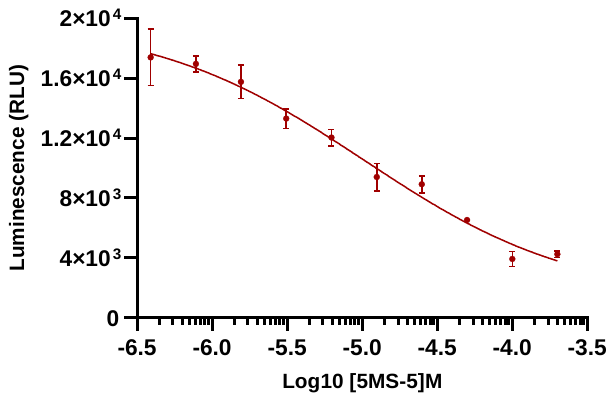

**Figure S2. Functional inhibition of CD28–B7 signaling by 5MS-5 in a bioluminescent reporter assay.** The CD28 Blockade Bioassay (Promega, Cat. #JA6101) was employed to assess the ability of **5MS-5** to inhibit CD28–B7 interactions in a co-culture system of CD28 Effector Cells (Jurkat) and aAPC/Raji Cells. The compound was evaluated in a 10-point dose–response format, and luminescence was quantified using the Bio-Glo™ Luciferase Assay System. Dose–response curve was fitted using nonlinear regression (four-parameter logistic model) in GraphPad Prism. Data are presented as mean ± SEM from n = 5 independent experiments.

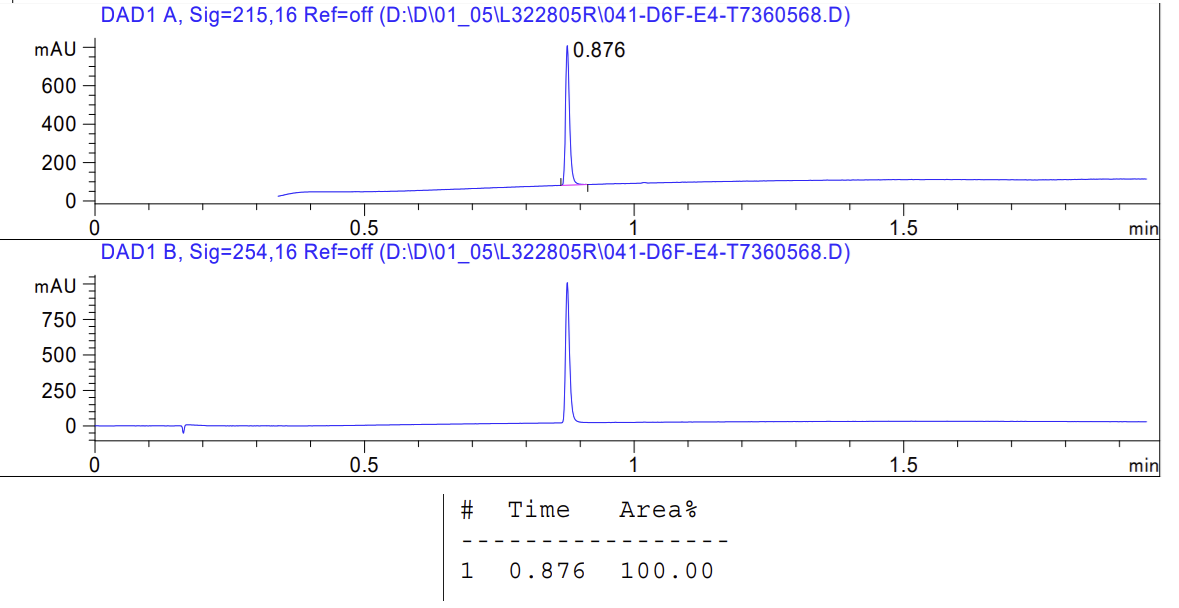

**Figure S3**. HPLC trace of compound **5MS**.

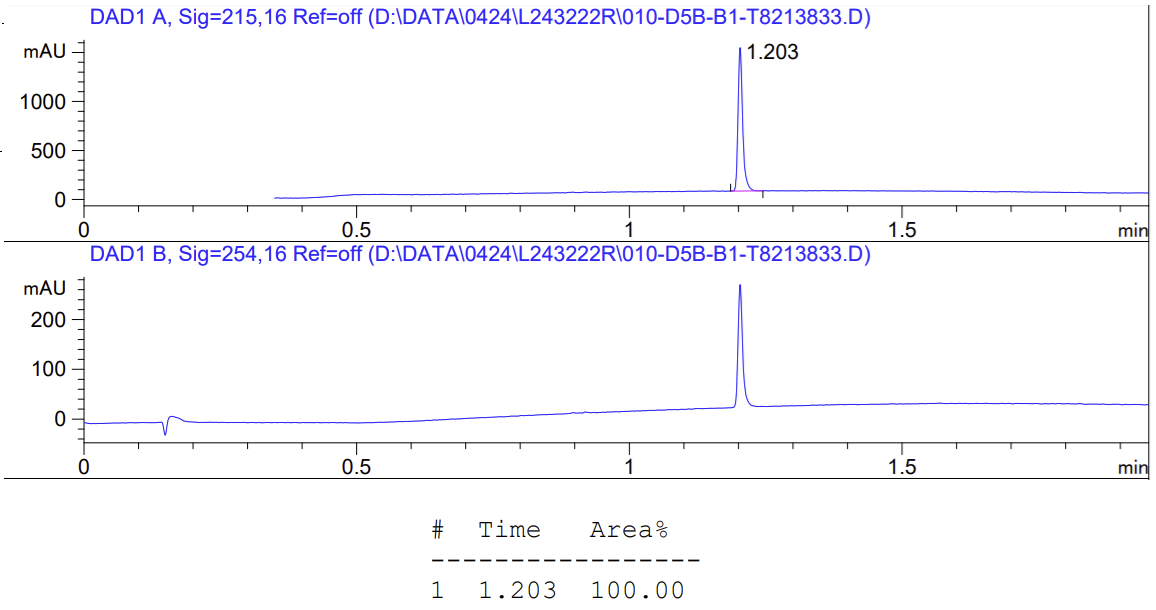

**Figure S4**. HPLC trace of compound **5MS-5**.

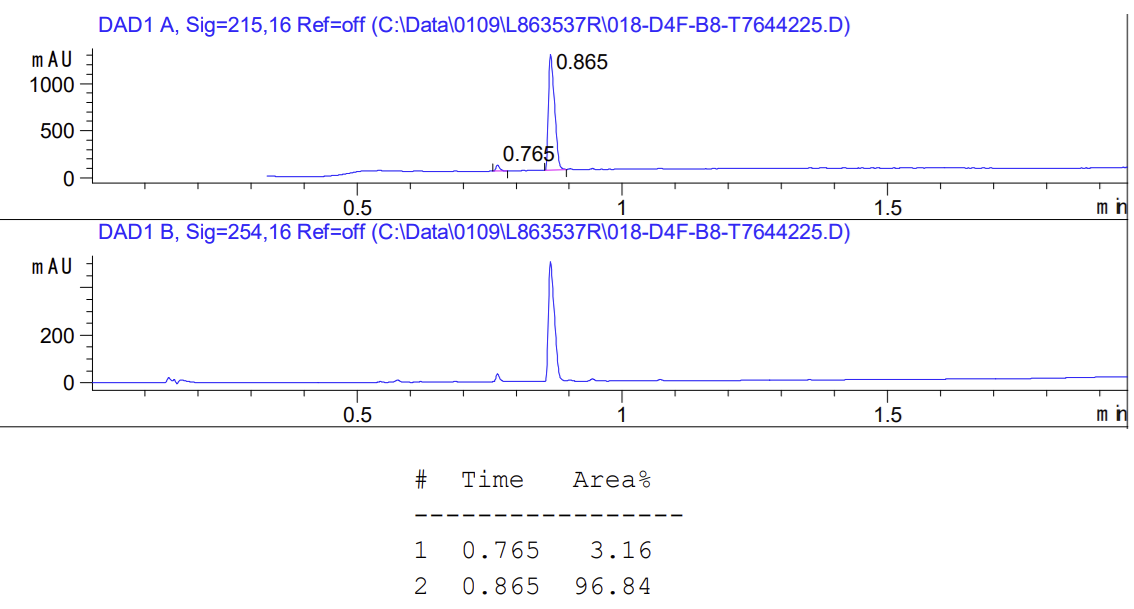

**Figure S5**. HPLC trace of compound **19MS**.

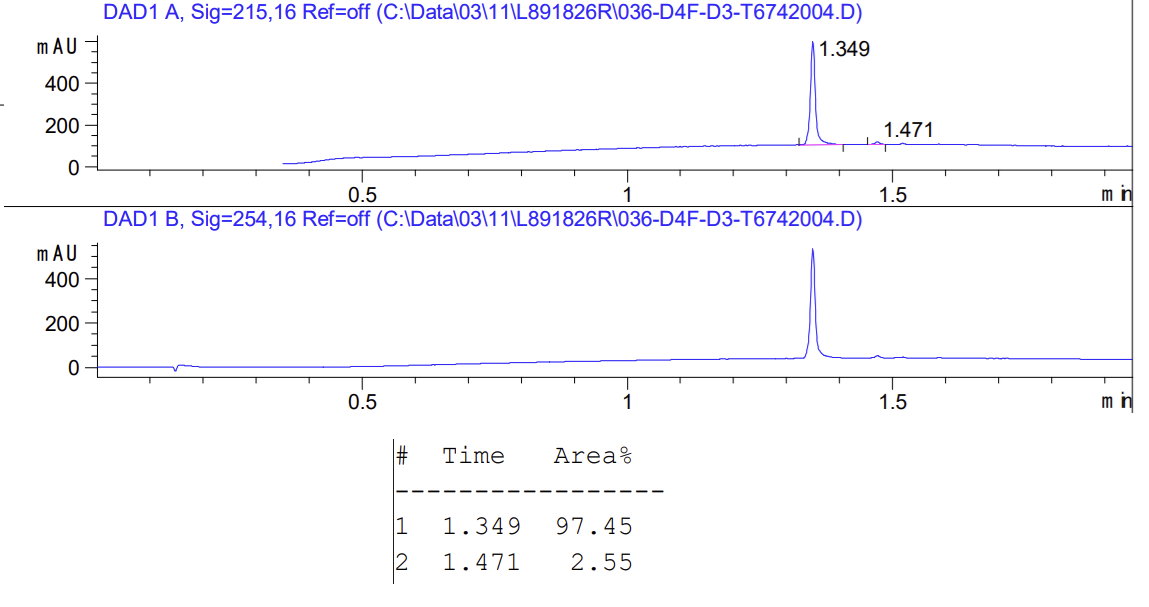

**Figure S6**. HPLC trace of compound **19MS-15**.
